## Supplementary Figures and Supporting Information for "*Trans* regulation of an odorant binding protein by a proto-Y chromosome affects male courtship in house fly"

**Supplementary Materials Summary**

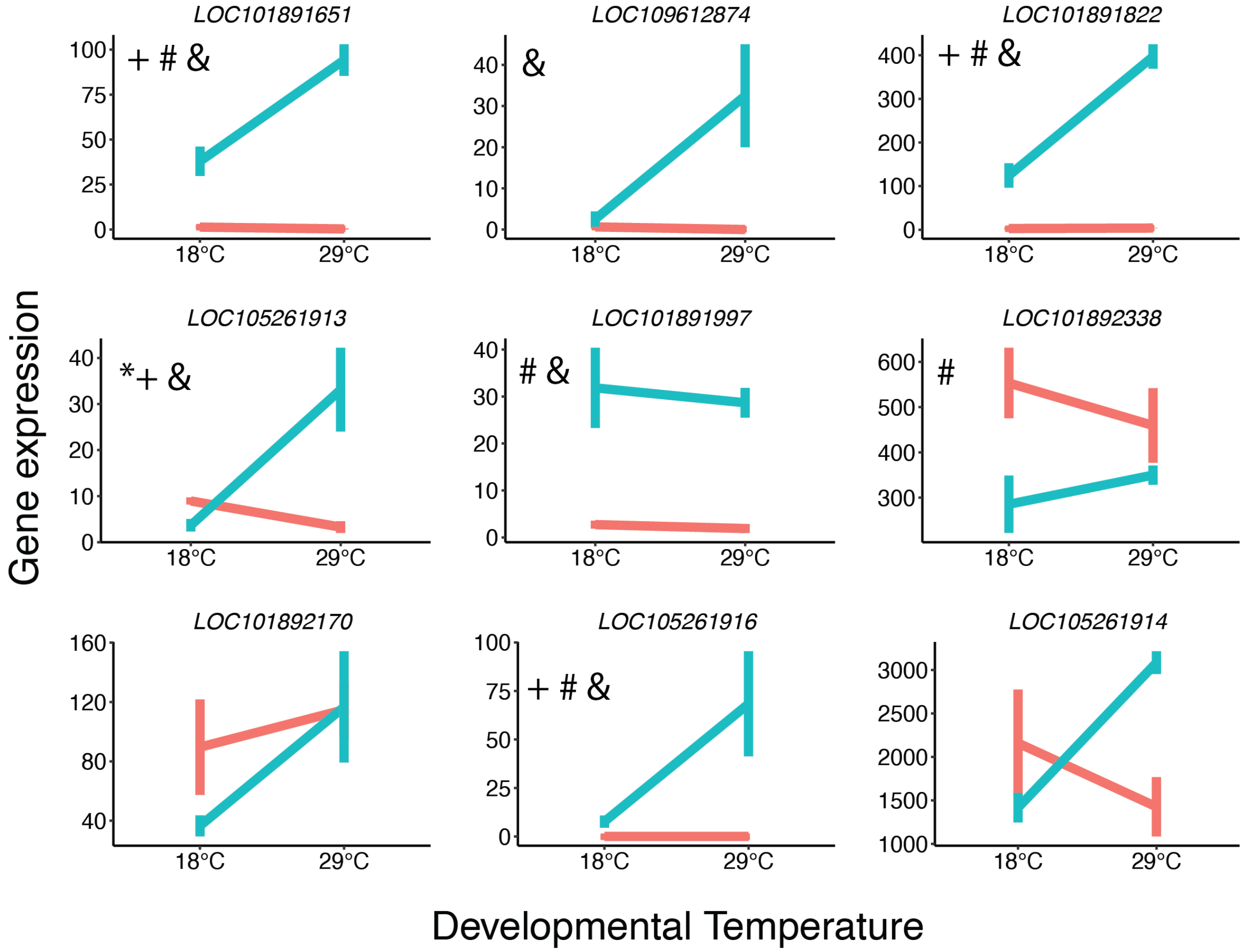


**Figure S1 -** Summary of *Obp56h* expression in $III^M$ (red) and $Y^M$ (blue) males reared at 18°C and 29°C. *Obp56h* genes are identified based on gene ID. Error bars denote standard errors of the mean. Significance of effects are shown by symbols: * significant G×T effect, +Temperature effect in YM males, # Genotype effect at 18°C, and & Genotype effect at 29°C. Data taken from Adhikari et al. (2021).

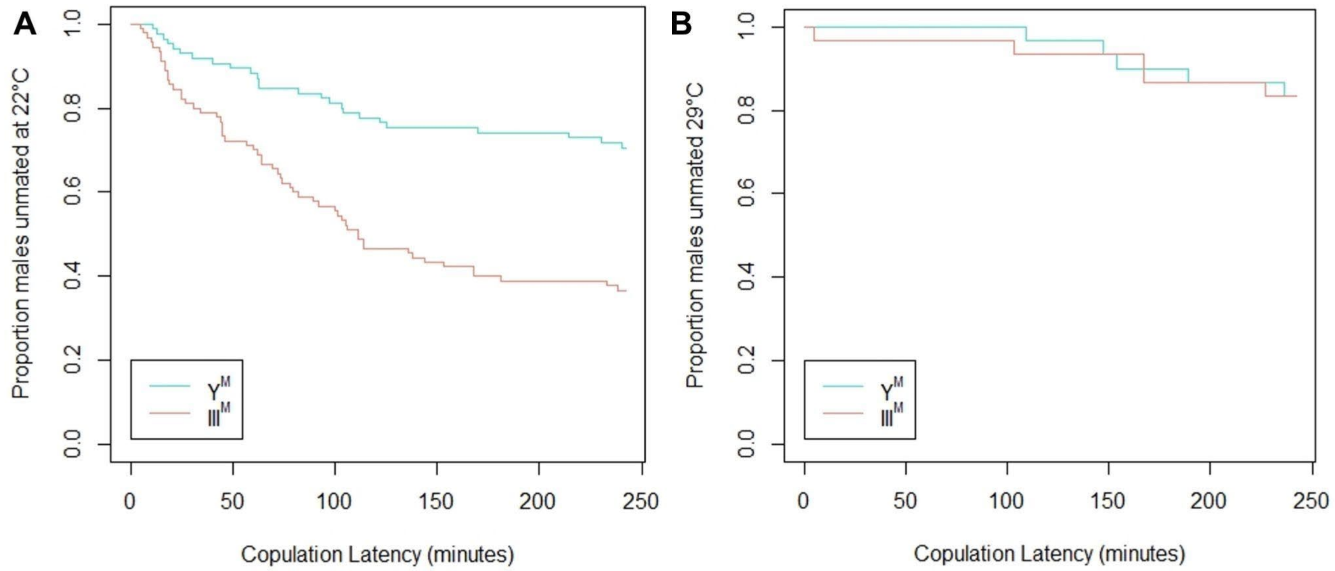


**Figure S2 -** Summary of survival analysis results (pooled across experimental batches) depicting the proportion of unmated Y$^{M}$ (turquoise) and III$^{M}$ (salmon) males as a function of time (minutes), at 22°C (panel A) and 29°C (panel B). Censored observations not depicted.

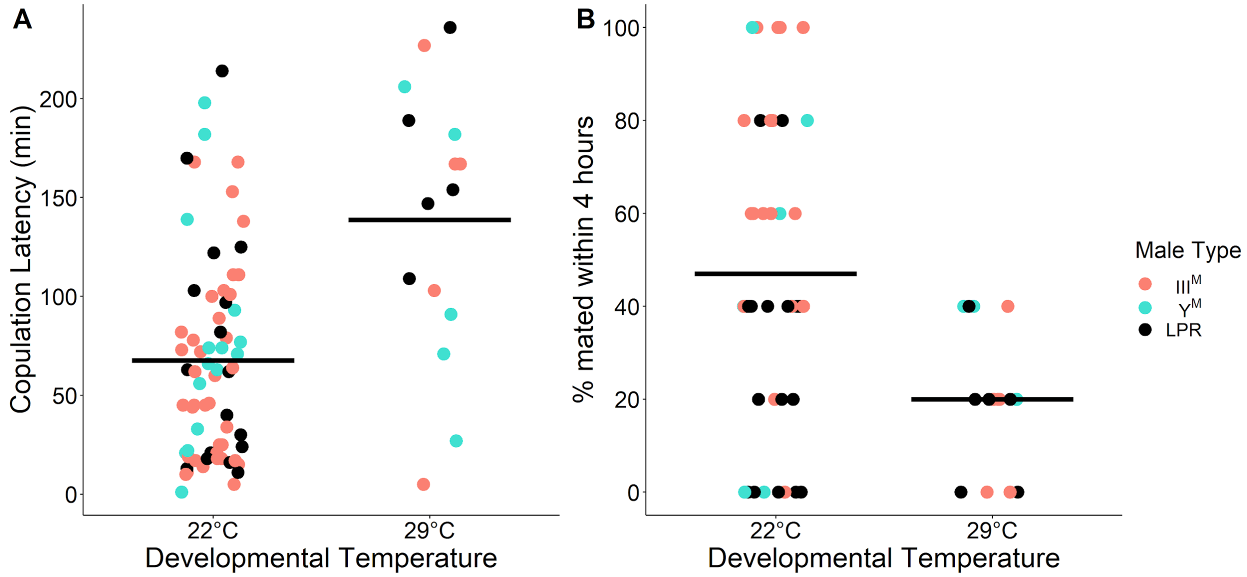


**Figure S3** - Summary of the effect of developmental temperature on copulation latency in III$^{M}$ (salmon), Y$^{M}$ (turquoise), and LPR (black) males reared at 22°C and 29°C. A) Copulation latency is estimated as the time taken to copulate (attached > 1 min) with a female. B) Copulation latency is estimated as a proportion of males (out of five males within one replicate) that mated with females within 4 hours within each experimental trial. All females used were from the LPR strain. Horizontal lines denote means within a given developmental temperature treatment.

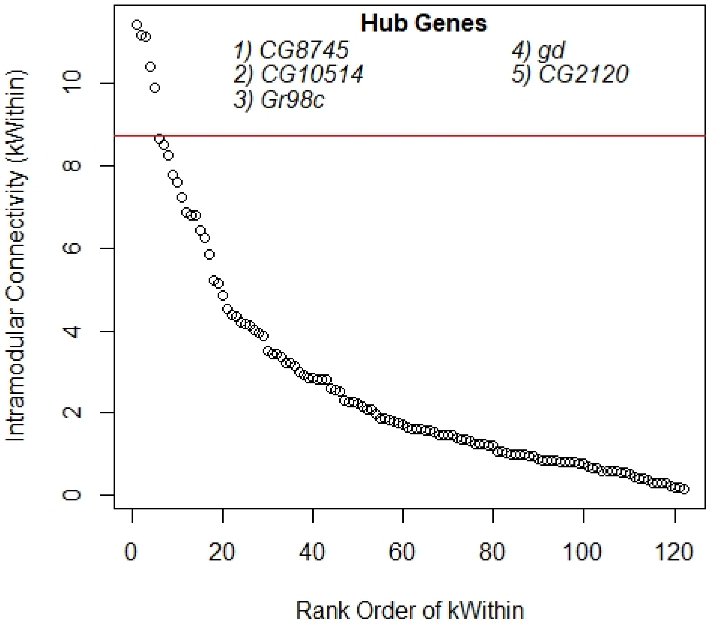


**Figure S4 -** Measures of intramodular connectivity (kWithin) for individual genes within the gene co-expression module, ranked in descending order. The horizontal red line denotes the separation between the top five hub genes (listed in descending order of kWithin) and the remainder of the genes in the module. Each data point indicates the kWithin value of an individual gene.

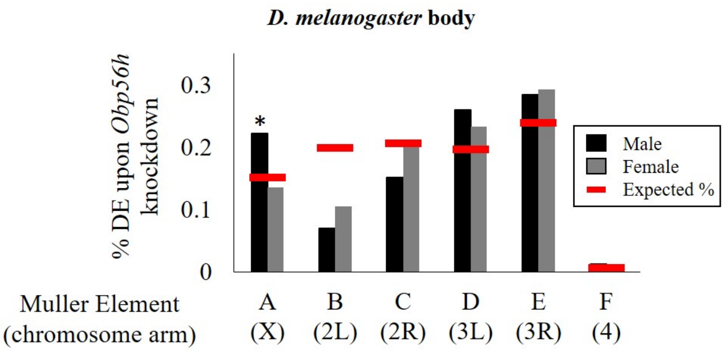


**Figure S5 -** Proportions of genes on each chromosome that are differentially expressed in the body (head removed) between *Obp56h* knockdown and control *D. melanogaster* (black bars: males, grey bars: females). Asterisks indicate a significant difference between observed (bars) and expected (red lines) counts of genes on each chromosome compared to all other chromosomes (Fisher's exact test, $p < 0.05$).

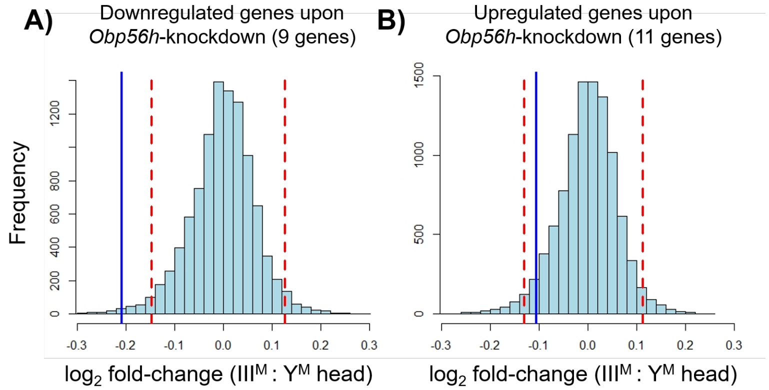


**Figure S6** - Differential expression between male house fly genotypes of genes that are differentially expressed upon *Obp56h* knockdown in *D. melanogaster*. Positive values indicate higher expression in $III^M$ house fly males, and negative values indicate higher expression in $Y^M$ males. The blue vertical line in each panel indicates the observed mean $\log_2$ fold-change for house fly transcripts of genes that are downregulated (A, 9 genes) or upregulated (B, 11 genes) upon *Obp56h* knockdown in *D. melanogaster* (Shorter et al. 2016). Histograms show the distributions of mean $\log_2$ fold-change values (A, for 9 randomly chosen genes; B, for 11 randomly chosen genes) drawn from a pool of all $\log_2$ fold- change differences in expression between $III^M$ and $Y^M$ males. Histograms were generated from 10,000 replicate draws of random genes without replacement. Dashed red lines show the boundaries for the middle 95% of the distributions.

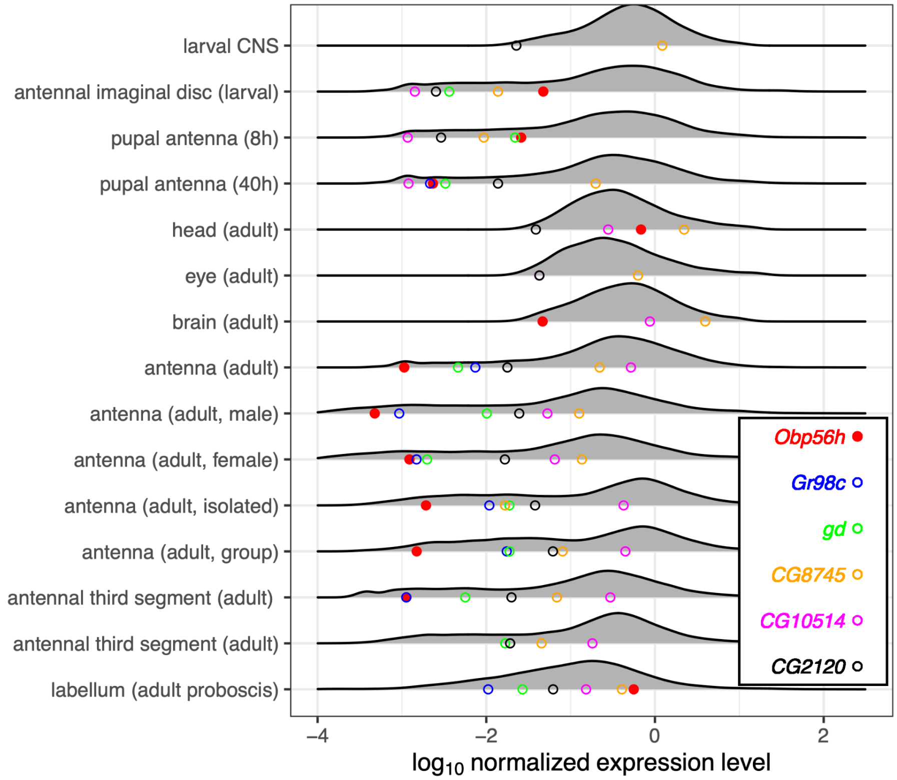

**Figure S7** - Expression levels in *D. melanogaster* head tissues of *Obp56h* and the hub genes in the co-expression network. The distribution of $\log_{10}$ normalized expression levels for all genes above a minimum threshold in a given tissue sample are plotted as smoothed histograms. Normalized expression was calculated for each gene or transcript in a tissue sample by dividing by the mean expression level across all genes or transcripts in the tissue sample. Tissue samples include two larval samples (central nervous system [CNS] and antennal imaginal disc), two pupal samples (antennae at 8 h post pupation and 40 h post pupation), and 11 adult samples. The expression levels of individual genes are shown as colored circles on the x-axis of each histogram. If a colored circle is not shown for a tissue, the gene corresponding to that color was not detected as expressed in the tissue.

### Supplementary Table Titles

**Table S1** - List of house fly samples used for RNA-seq analyses.
**Table S2** - List of *Drosophila melanogaster* samples used for tissue-specific expression analysis.
**Table S3 -** List of genes differentially expressed between $III^M$ and $Y^M$ male house fly head samples, according to limma (FDR < 0.05).
**Table S4 -** List of house fly genes, gene module assignments, and module membership values according to WGCNA.
**Table S5 -** Summary of gene ontology (GO) terms enriched within WGCNA module A.
**Table S6** - Summary of adjacency values and connection scores ($C_i$) for genes connected to *Obp56h* genes. The 100 genes with highest $C_i$ were classified as "central genes" with strong connections to *Obp56h* genes.
**Table S7** - List of gene ontology (GO) terms enriched within the lists of genes 1) identified as "central genes" with strong connections to *Obp56h* genes in male house flies, and 2) differentially expressed between *Obp56h*-suppressed and control male *D. melanogaster*, as reported in Shorter et al. (2016).
**Table S8** - Statistical tests comparing the fit of linear models predicting copulation latency between *CG2120* upregulated males and control male *D. melanogaster*.

### Supporting Information 1

*Summary of accumulated degree day (ADD) calculations for flies used in mating assays*

We used the following formula to estimate ADD of flies reared at different developmental temperatures for competitive and single-choice mating assays:

$$ADD = (T_D - T_t) \times d$$

where $T_D$ is the developmental temperature of a given fly, and $T_t$ is the threshold developmental temperature of 12.4°C, which was calculated for house flies in Wang et al (2018). Based on this value, male flies that developed at 18°C and 29°C for competitive mating assays had ADD ranges of 33.6-39.2 and 66.4-83 degree days (dd), respectively. While these ADD ranges do not overlap between developmental temperature treatments, these ranges represent time points during which adult house flies are mating in colony cages.

ADD male house flies developed at 18°C (competitive mating assays)

$$ADD = (18°C\text{-}12.4°C) \times (6d) = 33.6 \text{ dd}$$

$$ADD = (18°C\text{-}12.4°C) \times (7d) = 39.2 \text{ dd}$$

ADD male house flies developed at 29°C (competitive mating assays)

$$ADD = (29°C\text{-}12.4°C) \times (4d) = 66.4 \text{ dd}$$

$$ADD = (29°C\text{-}12.4°C) \times (5d) = 83 \text{ dd}$$

Male flies that developed at 22°C and 29°C in our single-choice mating assays had ADD ranges of 96-105.6 and 99.6-116.2 dd, respectively. Female flies used in these assays had ADD ranges of 100.8-113.4 dd. These ranges overlap with our estimate of ADD ranges for *Drosophila melanogaster* adults used in Shorter et al (2016): those flies had ADD ranges of 45-105 dd based on the reported age of assayed adults (3-7 d), and an estimated $T_t$ of 10°C (Loeb and Northrop 1917; Bliss 1926; Koštál et al. 2016).

ADD male house flies developed at 22°C (single-choice assays)

$$ADD = (22°C\text{-}12.4°C) \times (10d) = 96 \text{ dd}$$

$$ADD = (22°C\text{-}12.4°C) \times (11d) = 105.6 \text{ dd}$$

ADD male house flies developed at 29°C (single-choice assays)

$$ADD = (29°C\text{-}12.4°C) \times (6d) = 99.6 \text{ dd}$$

$$ADD = (29°C\text{-}12.4°C) \times (7d) = 116.2 \text{ dd}$$

ADD female house flies developed at 25°C (single-choice assays)

$$ADD = (25°C\text{-}12.4°C) \times (8d) = 100.8 \text{ dd}$$

$$ADD = (25°C\text{-}12.4°C) \times (9d) = 113.4 \text{ dd}$$

ADD fruit flies assayed in Shorter et al. 2016

$$ADD = (25°C\text{-}10°C) \times (3d) = 45 \text{ dd}$$

$$ADD = (25°C\text{-}10°C) \times (7d) = 105 \text{ dd}$$

### Supporting Information 2

*Summary of differential expression between $III^M$ and $Y^M$ males with a shared genetic background*

To account for the possibility that our main results might be influenced by the inclusion of $III^M$ males with a different genetic background than the rest of the sampled males, we conducted differential expression (DE) analysis on males with a shared (CS) genetic background, using the same pipeline as in the main study. These males with a shared genetic background differ only in their proto-Y chromosome ($Y^M$ or $III^M$). In this analysis, we identified 12 significantly DE genes between $III^M$ and $Y^M$ males. All of these genes are also DE in the full data set. The same *Obp56h* gene (*LOC105261916*) that was DE after Benjamini-Hochberg $p$ value adjustment in the full data set was also DE here. In addition, 5 more *Obp56h* genes had raw $p$ values $< 0.05$ in this data set, compared to 3 with raw $p < 0.05$ in the full data set. As in the full data set, we find a similar trend of enrichment of Muller Element A genes within this list of DE genes (4/12 DE genes assigned to Muller Element A). However, this was not statistically significant, possibly due to the small number of DE genes (Fisher's exact test $p = 0.060$). Due to low power from the smaller number of RNA-seq samples in this data set, we did not perform WGCNA.

**Supporting Information 3**

*Discussion of allele-specific expression results*

Previous studies have shown that the $III^M$ and $Y^M$ nearly isogenic lines included in our allele-specific expression analysis are minimally genetically diverged (Meisel et al. 2017; Son and Meisel 2021), making it difficult to identify fixed diagnostic sites that are consistent across $III^M$ strains. We obtained data across three studies (Meisel et al. 2015; Son et al. 2019; Adhikari et al. 2021) that sampled two $III^M$ strains (CS and CSrab). We did not require CS and CSrab males to share the same $III^M$ alleles because they carry different naturally derived $III^M$ chromosomes, and there is a paucity of fixed $III^M$ alleles. Instead, we only required that SNP alleles were consistent across experiments within each strain. In the case of *LOC101893264* (*Md-gd*), at all seven diagnostic sites, all reads mapped to either the reference or alternate allele, and allele assignments were consistent within strains. However, the specific exonic alleles differ between the two strains because there are very few fixed differences between $III^M$ and III. Although CS and CSrab males possess different alleles within *Md-gd* exons, they may ultimately possess the same *cis*-regulatory allele for this gene—an allele which differs from that of $Y^M$ males. That shared *cis*-regulatory allele would be responsible for up-regulation of the $III^M$ allele in both CS and CSrab males. In $Y^M$ males, conversely, only one read was mapped to any of the diagnostic sites within *Md-gd*, suggesting that *Md-gd* is not expressed from the standard chromosome III in the tissues and conditions we sampled.

In the case of *LOC101893651* (orthologous to *CG2120*), all diagnostic SNP sites in exonic regions are consistent both across experiments and across $III^M$ strains, with all reads mapping to the non-reference allele at each site. In $Y^M$ males, a total of three reads were mapped across all diagnostic sites. Together, this suggests that CS and CSrab $III^M$ males are either homozygous for the non-reference allele (unlikely because they are $III^M$/III heterozygotes), or they exhibit monoallelic gene expression in this gene.

In the case of *LOC101894501* (homologous to *Gr98c*), all diagnostic SNP sites in exonic regions are consistent both across experiments and across $III^M$ strains. In $III^M$ males, a total of 65 reads were assigned to the reference (III) allele, and 169 reads were assigned to the non-reference ($III^M$) allele. In $Y^M$ males, a total of four reads were mapped across all diagnostic sites, all of which were assigned to the reference (standard chromosome III) allele. Together, these results suggest that CS and CSrab III$^{M}$ males exhibit III$^{M}$-biased, but not monoallelic, gene expression. However, the higher expression of the III allele in III$^{M}$ males than Y$^{M}$ males suggests that *trans* regulators further increase the expression of *Md-Gr35* in III$^{M}$ males. Therefore, we conclude that *Md-Gr35* is likely differentially regulated by a combination of *cis* and *trans* regulatory elements.

**Supporting Information 4**

*Expression of Obp56h and co-expressed genes*

To further evaluate the relationships between *Obp56h* and the hub genes in the co-expression network, we investigated their expression across *D. melanogaster* head tissues (Fig. S7). We confirmed that *Obp56h* is highly expressed in adult head, and we found that expression is most pronounced in the labellum (i.e., proboscis). While *Obp56h* has undetectable expression in the adult antennae, it is expressed in the larval imaginal disc destined to develop into antenna and eye tissues, as well as the developing antennae of early pupae. This suggests that if *Obp56h* functions in adult antennae, it must either persist via developmental expression or be transported there from another part of the head or body. In general, these expression patterns are consistent with *Obp56h* involvement in chemosensation, either via taste or smell.

The labellum is the only head component in which all five hub genes are expressed in *D. melanogaster* (Fig. 7). Notably, both *Obp56h* and *Gr98c* are expressed in labellum (Fig. S7), consistent with the co-regulation we propose above. In contrast, *gd* is expressed in pupal antennae, along with *Obp56h* (Fig. S7). It is therefore possible that the regulatory connection between *gd* and *Obp56h* that we propose occurs in the developing antennae. Two of the other hub genes are highly expressed in most or all head tissue samples (Fig. S7). *CG8745* is highly or moderately expressed across developing and adult head tissues. *CG10514* is also highly expressed across adult head tissues (including antennae), although not in the larval or pupal antennae.
